## Supporting Information for "Computer-guided Binding Mode Identification and Affinity Improvement of an LRR Protein Binder without Structure Determination"

Hak-Sung Kim

Department of Biological Sciences, Korea Advanced Institute of Science and Technology, Daejeon, 34141, Republic of Korea.

**Figure S1. The repebody structure.** (**A**) A repebody (Rb) largely consists of three parts: N-termianl cap (LRRNT), variable regions (LRRV) and C-terminal cap (LRRCT). Binding occurs at the concave region of LRRV (in darker blue). (**B**) Structure of a single LRRV motif, with side chains of conserved residues rendered as stick figures. Each LRR is composed of six conserved leucine residues, a central conserved asparagine residue, and conserved phenylalanine residue on the C-terminal side.

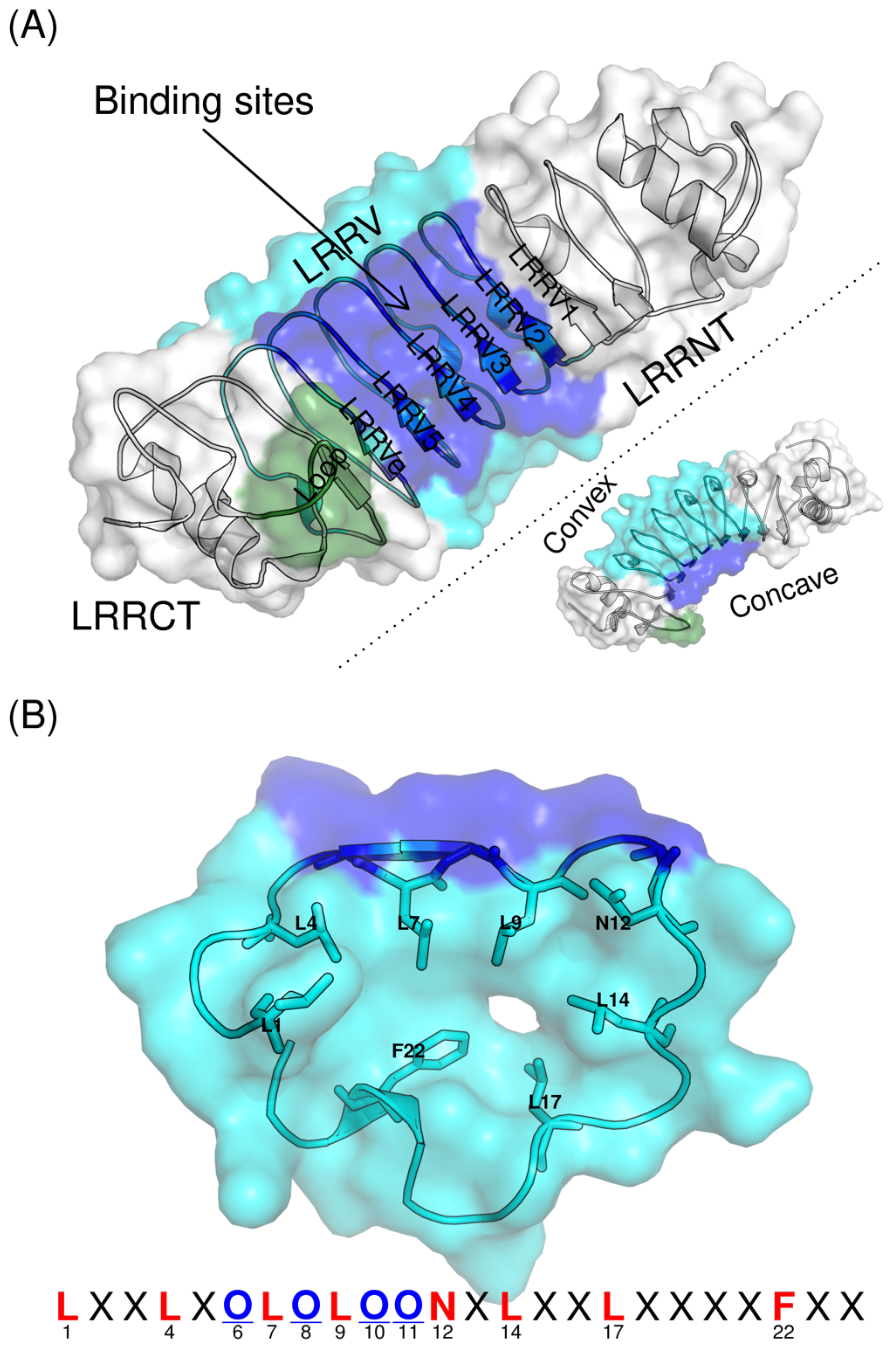

**Figure S2. Detailed information of the mutations for hFc-F4 epitope localization and titration curves.** Based on the $K_d$ values, H310 and N315 overlap epitopes.

| | | Mutations | N | $K_d$ (nM) | ΔH (cal/mol) | ΔS (cal/mol/deg) | ΔG (cal/mol) |
|---|---|---|---|---|---|---|---|
| Wild Type | | | 1.11 ± 0.00692 | 128.7 ± 18.59 | –11130 ± 109.8 | –5.79 | –9409.5 ± 109.8 |
| Var 1 | | Q362E N389E N390K | 0.93 ± 0.00695 | 119.6 ± 20.77 | –8621 ± 104.5 | 2.76 | –9441.1 ± 104.5 |
| Var 2 | | H268K E269K R292L | 0.93 ± 0.00461 | 121.5 ± 13.74 | –9352 ± 76.0 | 0.28 | –9435.2 ± 76.0 |
| Var 3 | | H310A N315K H435K | 1.01 ± 0.00124 | 408.2 ± 67.20 | –6073 ± 106.2 | 8.87 | –8708.7 ± 106.2 |
| Var 3 | H310A | H310A | 1.01 ± 0.00542 | 194.9 ± 19.46 | –8739 ± 71.8 | 1.39 | –9152.0 ± 71.8 |
| | N315K | N315K | 0.96 ± 0.00794 | 187.3 ± 29.15 | –8490 ± 108.1 | 2.31 | –9176.4 ± 108.1 |
| | H435K | H435K | 1.04 ± 0.00532 | 98.0 ± 13.41 | –8955 ± 77.4 | 2.04 | –9561.2 ± 77.4 |

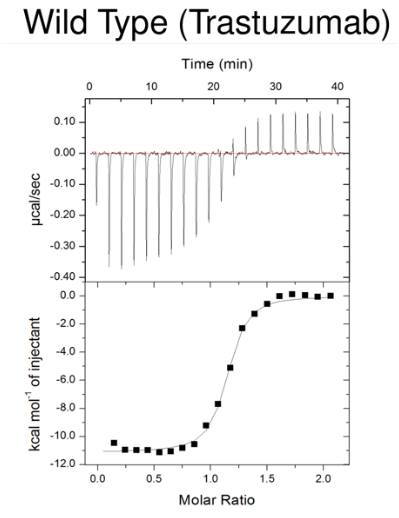

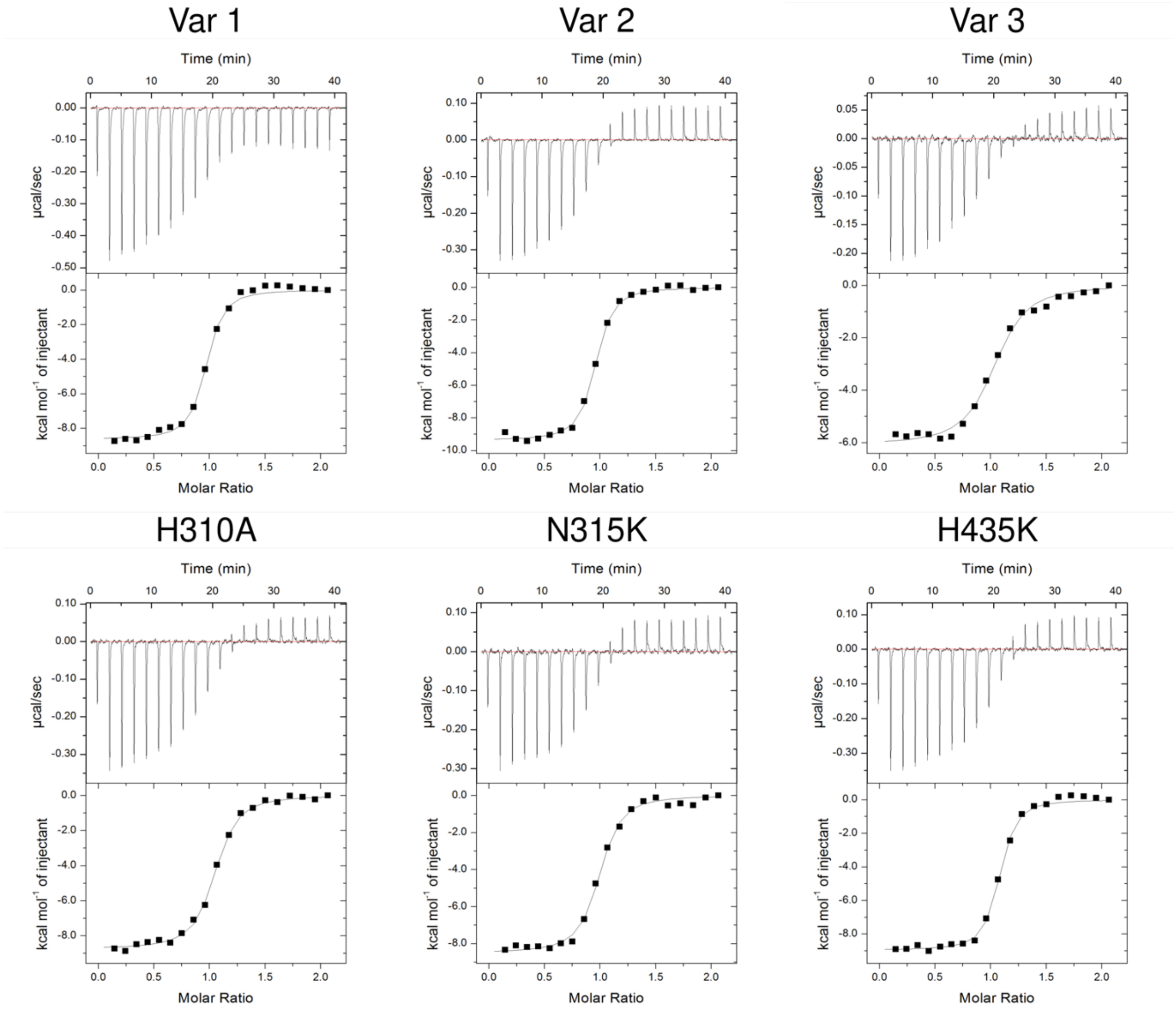

**Figure S3. Docking models of RbF4.** The crystal structure is in gold. (A) The full atom energy minimization step may change the overall structure, but the binding interactions are likely to be maintained. The I-RMSD value of the lowest energy model (Model 1, blue) is slightly higher than the model with the second lowest energy (Model 2, pink). However, its $f_{nat}$ is twice higher. (B) There are two docking models which are in contact with the three mutations in Var 3 (Model 9: cyan and Model 10: pink). While their binding interface regions are largely correct, the binding orientation of the lower energy model (pink) is completely inverted.

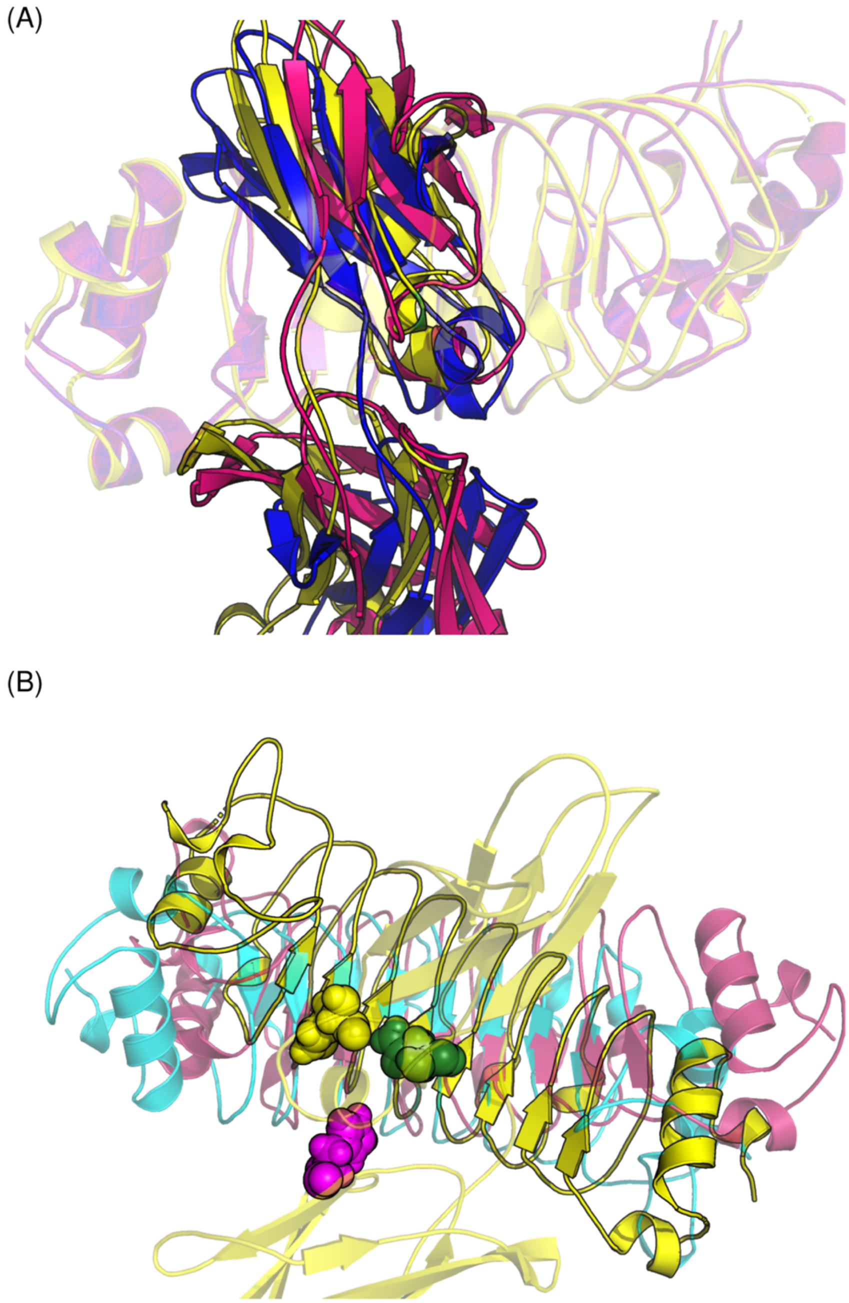

**Figure S4. Details of binding affinities of RbF4 variants (RbF4, RbF4-LT and RbF4-MR) to $hIgG_1$, $mIgG_1$ and rIgG.** RbF4 binds strongly to $hIgG_1$ and weakly to m $IgG_1$. However, no binding affinity is measured for RbF4-rIgG. The truncation of the loop (RbF4-LT) enabled the variant to bind to all IgGs with similar binding affinities. RbF4-MR gains strong binding affinities for $hIgG_1$ and $mIgG_1$. See **Supplementary Table 3** for details.

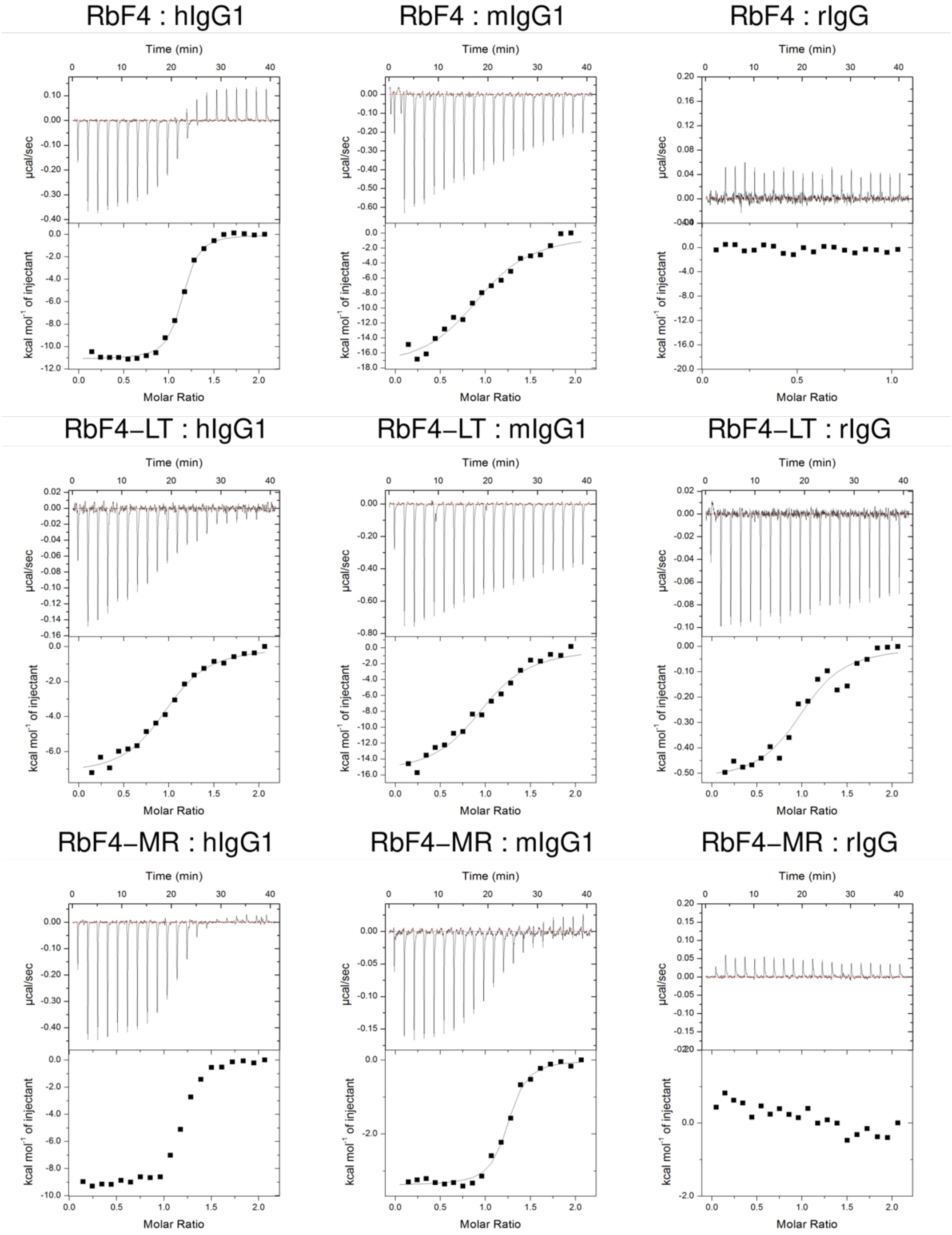

**Figure S5. FoldX scan of the RbF4 loop.** The residue scan using FoldX suggests that the inclusion of the loop may not enhance the binding affinity of RbF4 for rIgG.

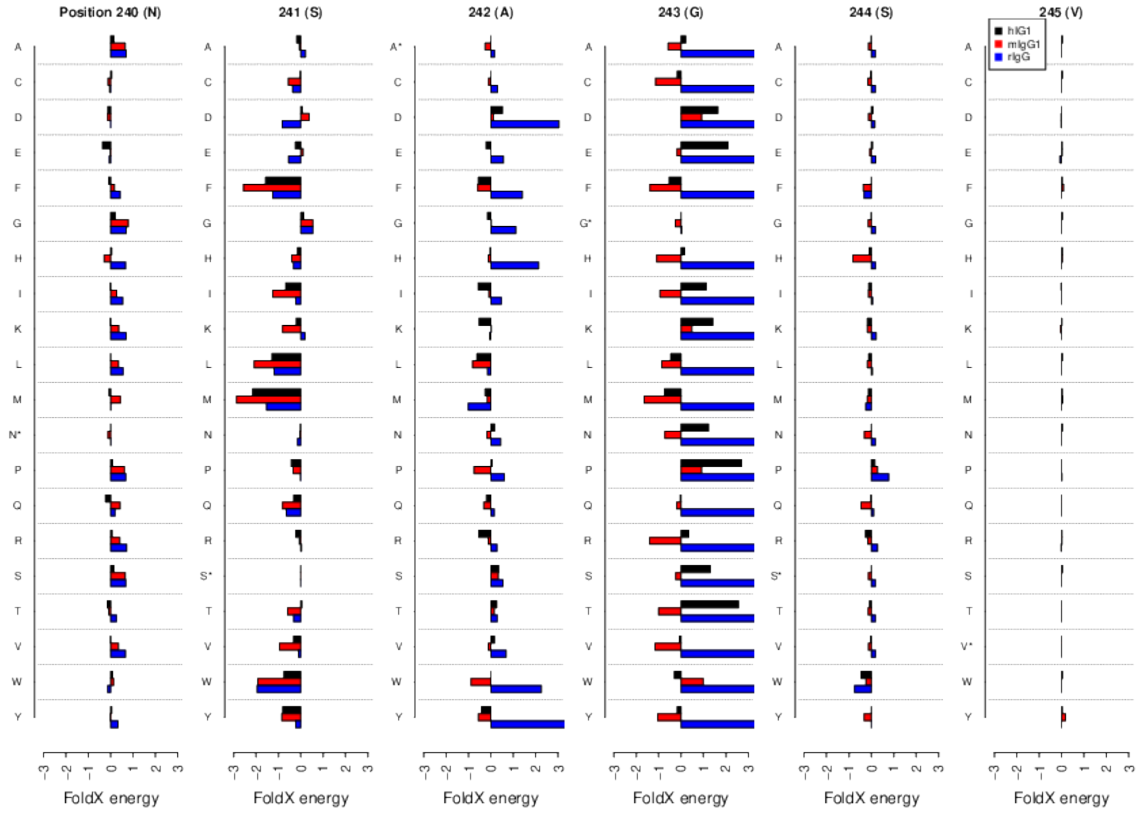

**Figure S6. Predicted ΔΔG values of the RbF4 variants.** The variant with S241M and S244R mutations (RbF4-MR) is predicted to bind to both $hIgG_1$ and $mIgG_1$ with strong binding affinities. S244C and S244P were not considered.

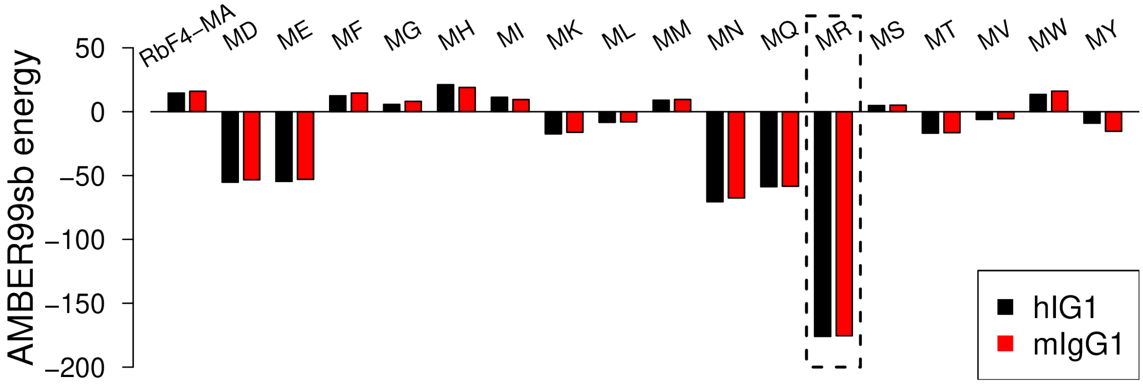

**Figure S7. Binding mode prediction with the ClusPro score.** Docking models that are in contact with epitope overlapping residues (localized docking models) are in solid circle. The blue circle is the model with the lowest ClusPro score. Crystal structures are in yellow and the docking models with the lowest energies are in blue on the right hand side. Score assessment using the ClusPro score is not predictive.

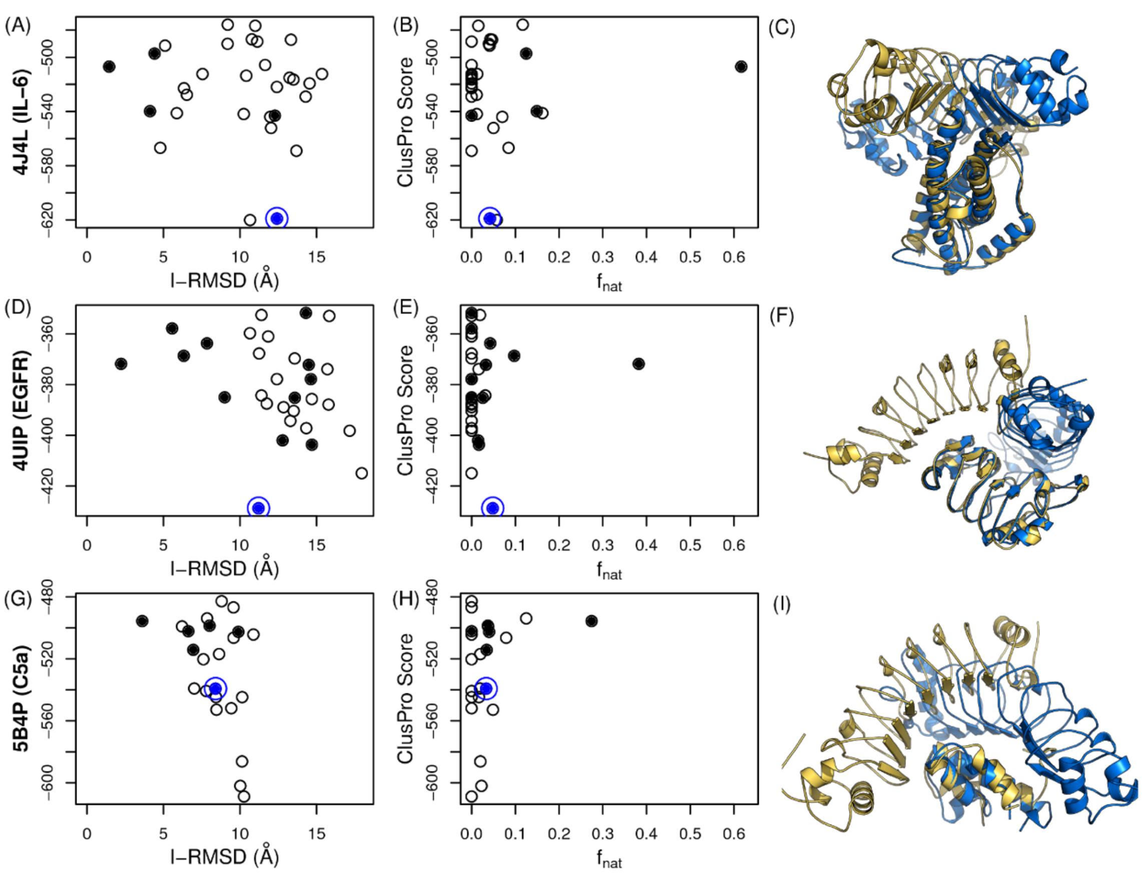

**Table S1. Data collection and refinement statistics**

| | RbF4/hFc |
|---|---|
| **Data collection** | |
| Space group | P $2_12_12_1$ |
| Cell dimensions | |
| *a*, *b*, *c* (Å) | 59.9, 107.4, 171.4 |
| $\alpha$, $\beta$, $\gamma$ (°) | 90,90,90 |
| Resolution (Å) | 30-3.0 (3.05-3.00)* |
| $R_{merge}$ (%) | 17.5(100.0) |
| $I/\sigma I$ | 15.0(1.7) |
| Completeness (%) | 100(100) |
| Redundancy | 13.2(12.6) |
| **Refinement** | |
| Resolution (Å) | 30-3.0 |
| No. reflections | 22835 |
| $R_{working}$ / $R_{free}$ (%) | 25.7 / 33.3 |
| No. atoms | |
| Protein | 6613 |
| Ligand/ion | 99 |
| water | 6 |
| B-factors | |
| Protein | 56.9 |
| Ligand/ion | 76.4 |
| water | 35.2 |
| R.m.s deviations | |
| Bond lengths (Å) | 0.010 |
| Bond angles (º) | 1.246 |

$I/\sigma I$, mean intensity/sigma of all reflections; $R_{free}$, $\Sigma||F_{obs}|-|F_{calc}|| / \Sigma|F_{obs}|$ where 5% of randomly selected data were used; $R_{merge}$, $\Sigma|I_{hkl}-<I_{hkl}>| / \Sigma I_{hkl}$ ; where $<I_{hkl}>$ is the mean intensity of all reflections equivalent to refection hkl; $R_{work}$, $\Sigma||F_{obs}|-|F_{calc}|| / \Sigma|F_{obs}|$.

*Highest resolution shell is shown in parenthesis.

**Table S2. Quality measures and contact information of RbF4 docking models.** Each black block indicates that the Fc position is in contact with the docked repebody.

| **Model** | **AMBER99sb total energy** | **I-RMSD** | **$f_{nat}$** | **Var 1** | | | **Var 2** | | | **Var 3** | | |
|---|---|---|---|---|---|---|---|---|---|---|---|---|
| | | | | **Q362** | **N389** | **N390** | **H268** | **E269** | **R292** | **H310** | **N315** | **H435** |
| **1** | -23025.43 | 2.886 | 0.434 | | | | | | | ■ | ■ | |
| **2** | -23002.86 | 2.323 | 0.292 | | | | | | | ■ | ■ | |
| **3** | -22984.81 | 14.954 | 0.044 | | | | | | | ■ | ■ | |
| **4** | -22981.08 | 15.629 | 0.057 | | | | | | | ■ | ■ | |
| **5** | -22978.22 | 15.264 | 0.016 | | | | | | | ■ | ■ | |
| **6** | -22964.96 | 4.057 | 0.041 | | | | | | | ■ | ■ | |
| **7** | -22957.28 | 6.922 | 0.000 | | | | | | | ■ | ■ | |
| **8** | -22997.36 | 16.006 | 0.056 | | | | | | | ■ | ■ | ■ |
| **9** | -22990.97 | 3.901 | 0.279 | | | | | | | ■ | ■ | ■ |
| **10** | -23033.53 | 14.573 | 0.000 | | | | ■ | ■ | ■ | | | |
| **11** | -23012.27 | 14.588 | 0.046 | | ■ | ■ | | | ■ | ■ | | |
| **12** | -23010.70 | 14.071 | 0.046 | | ■ | ■ | | | ■ | ■ | | |
| **13** | -23010.70 | 13.929 | 0.000 | | | | | | ■ | | | |
| **14** | -22997.03 | 15.852 | 0.015 | | | | ■ | ■ | | | | |
| **15** | -22994.80 | 12.450 | 0.000 | | ■ | | | | | | | ■ |
| **16** | -22991.84 | 11.205 | 0.000 | | | | ■ | ■ | | | | |
| **17** | -22989.76 | 15.473 | 0.017 | | | | | | ■ | | | |
| **18** | -22984.90 | 19.531 | 0.000 | | | | | | | | | |
| **19** | -22983.64 | 12.852 | 0.000 | | ■ | | | | | | | ■ |
| **20** | -22978.22 | 15.716 | 0.000 | | | | ■ | ■ | ■ | | | |
| **21** | -22976.98 | 15.060 | 0.000 | | | | ■ | ■ | ■ | | | |
| **22** | -22976.54 | 15.383 | 0.000 | | | ■ | ■ | ■ | ■ | | | |
| **23** | -22973.94 | 8.051 | 0.000 | | | | | | | ■ | | |
| **24** | -22970.76 | 18.285 | 0.017 | | | | ■ | ■ | | | | |
| **25** | -22965.44 | 14.074 | 0.000 | | | | | | | | | |
| **26** | -22957.28 | 10.174 | 0.000 | | | | | | ■ | ■ | | |
| **27** | -22948.56 | 12.976 | 0.000 | | | | ■ | ■ | ■ | | | |
| **28** | -22945.15 | 14.537 | 0.000 | | | | ■ | ■ | ■ | | | |
| **29** | -22936.86 | 15.529 | 0.000 | | | | | | | | | |

**Table S3. In silico binding specificity control and affinity improvement.** RbF4 strongly binds to $hIgG_1$ (weakly to $mIgG_1$), but no binding to rIgG is observed. Removal of the loop (RbF4-LT) causes a slight reduction in the binding affinity toward $hIgG_1$, but yields a marginal affinity improvement for $mIgG_1$. RbF4-LT produces a significant improvement in the affinity for rIgG. RbF4 with two mutations at 241 and 244 (S241M with S244R, RbF4-MR) binds to both $hIgG_1$ and $mIgG_1$ with high binding affinities.

| **Target** | **$hIgG_1$** | | | **$mIgG_1$** | | | **rIgG** | | |
|---|---|---|---|---|---|---|---|---|---|
| **Rb** | **RbF4** | **RbF4-LT** | **RbF4-MR** | **RbF4** | **RbF4-LT** | **RbF4-MR** | **RbF4** | **RbF4-LT** | **RbF4-MR** |
| **$K_d$ (nM)** | 128.70 | 598.80 | 125.79 | 1009.08 | 775.19 | 168.07 | N. D. | 1189.06 | > 6 μM |
| **N** | 1.01 ± 0.0092 | 0.984 ± 0.0190 | 1.19 ± 0.0078 | 0.964 ± 0.0356 | 0.989 ± 0.0326 | 1.21 ± 0.0099 | 7.83e4 ± 5.03e12 | 1.01 ± 0.0455 | 1.87 ±0.173 |
| **ΔH (cal/mol)** | -8102 ±110.7 | -7335 ±198.9 | -9394 ±100.5 | -2.1e4 ±1139 | -1.593e4 ± 752.6 | -3386 ±43.58 | -325.8 ±162.7 | -529.9 ±33.4 | 2107 ±341.9 |
| **ΔS (cal/mol/deg)** | 2.65 | 3.9 | 0.08 | -43.6 | -25.5 | 19.6 | 36.3 | 25.3 | 30.7 |
